# Deciphering the Evolutionary History of Complex Rearrangements in Head and Neck Cancer Patients Using Multi-Omic Approach

**DOI:** 10.1101/2022.08.19.504509

**Authors:** Jeesoo Chae, Jin Seok Lee, Jongkeun Park, Dong-Sung Lee, Weon Seo Park, Benjamin Clock, Jesse R. Dixon, Yuh-Seog Jung, Dongwan Hong

## Abstract

Despite the large efforts in international cancer genome consortium studies, there are still a large proportion of tumors with complex genomic rearrangement often remained without a clinically relevant molecular characterization. Integration of multi-omic data helps elucidating evolutionary history of such cases and identifying predictive molecular markers. Here we present the findings of our proof-of-principle study that investigated the evolutionary history of complex rearrangements in primary head and neck tumor genomes integrating long-read whole-genome, Hi-C, and RNA sequencing. We report a HPV-positive case with development of complex genomic rearrangements tracing back to HPV-mediated genomic instability and a HPV-negative case with an enhancer hi-jacking in a region of chromothripsis predicted to co-occur with a neoloop and a super-enhancer. These structural alterations resulted in overexpression of the oncogenes *CCND1* and *ALK*, respectively, validated with immunohistochemistry assay. Furthermore, we introduce a novel analytic approach utilizing long-read whole-genome data distinguishing somatic mutations before and after structural variants. Our findings highlight the need for multi-modal sequencing strategies to increase our understanding of cancer evolution and rare biomarkers in poorly understood cancers.

## INTRODUCTION

Somatic structural variants (SVs) are a class of large variants that bear remarkable implications in the evolutionary history of malignancies (Lee *et al*, 2019). Recent large-scale studies have extensively profiled tumor genomes using whole-genome sequencing (WGS) and revealed driver SVs responsible for oncogenesis across various types of primary tumors (Consortium, 2020; Gerstung *et al*, 2020; Jamal-Hanjani *et al*, 2017; Mitchell *et al*, 2018). However, still a considerable proportion of tumors with complex genome rearrangements are rarely cataloged systematically and frequently remain neglected. One of the reasons of this is due to the technical shortage of the popularly used short-read (SR) WGS which leads to high false positive rates for SV detection (Mahmoud *et al*, 2019), and another reason is the lack of complementary knowledge to approach the tumor genome from various angles.

Long-read (LR) high-fidelity (HiFi) circular consensus sequencing (CCS) provides unique opportunities to call SVs at a higher sensitivity rate and at single-molecule resolution owing to its read lengths that span tens of kilobases long with low base error rates (Wenger *et al*, 2019). To date, HiFi CCS has only been used to study two breast cancer patients’ genomes (Aganezov *et al*, 2020). SVs, especially translocations, accompany conformational changes to the three-dimensional (3D) genome organization (Spielmann *et al*, 2018), which can be detected using Hi-C sequencing. Integration of the two sequencing modalities therefore increases SV detection performance.

Here, we report the results of analysis of highly annotated somatic structural variants in human papillomavirus (HPV) positive and negative primary tumors of the head and neck. We employed a novel approach to infer the timing of somatic mutations emergence and reconstructed evolutionary history of structural changes through integrative analysis of multilayered data, including *de novo* assembly and neo-loop prediction analysis, which eventually ended up with disruption of oncogenes. For this, we performed SR and LR (HiFi CCS) WGS, Hi-C sequencing, and RNA sequencing (RNA-seq). This is the first study to comprehensively combine and analyze data generated by the above sequencing modalities of the same tumor samples derived from head and neck cancer (HNC) patients.

## RESULTS AND DISCUSSION

To investigate the oncogenic role of genomic rearrangements in tumorigenesis, we selected one HPV-positive (HNT-140) and one HPV-negative (HNT-073) primary HNC patients whose tumor genomes did not harbor driver single-base substitutions (SBSs) but exhibited substantial SV burden from an internal cohort of HNC patients (Fig 1A). The clinicopathological data and generated sequencing data statistics are summarized in Table EV1 and EV2, respectively. After comprehensive correction of the SV breakpoints in both LR and SR WGS due to the limited accuracy of existing SV detection algorithms, we found a total of 134 and 598 somatic SVs in HNT-140 and HNT-073, respectively, across SR and LR callsets (Fig EV1, and Tables EV2-EV6). Compared to SR WGS, LR WGS identified more insertions and yielded less variable breakpoints (Fig EV1).

**Figure 1.**
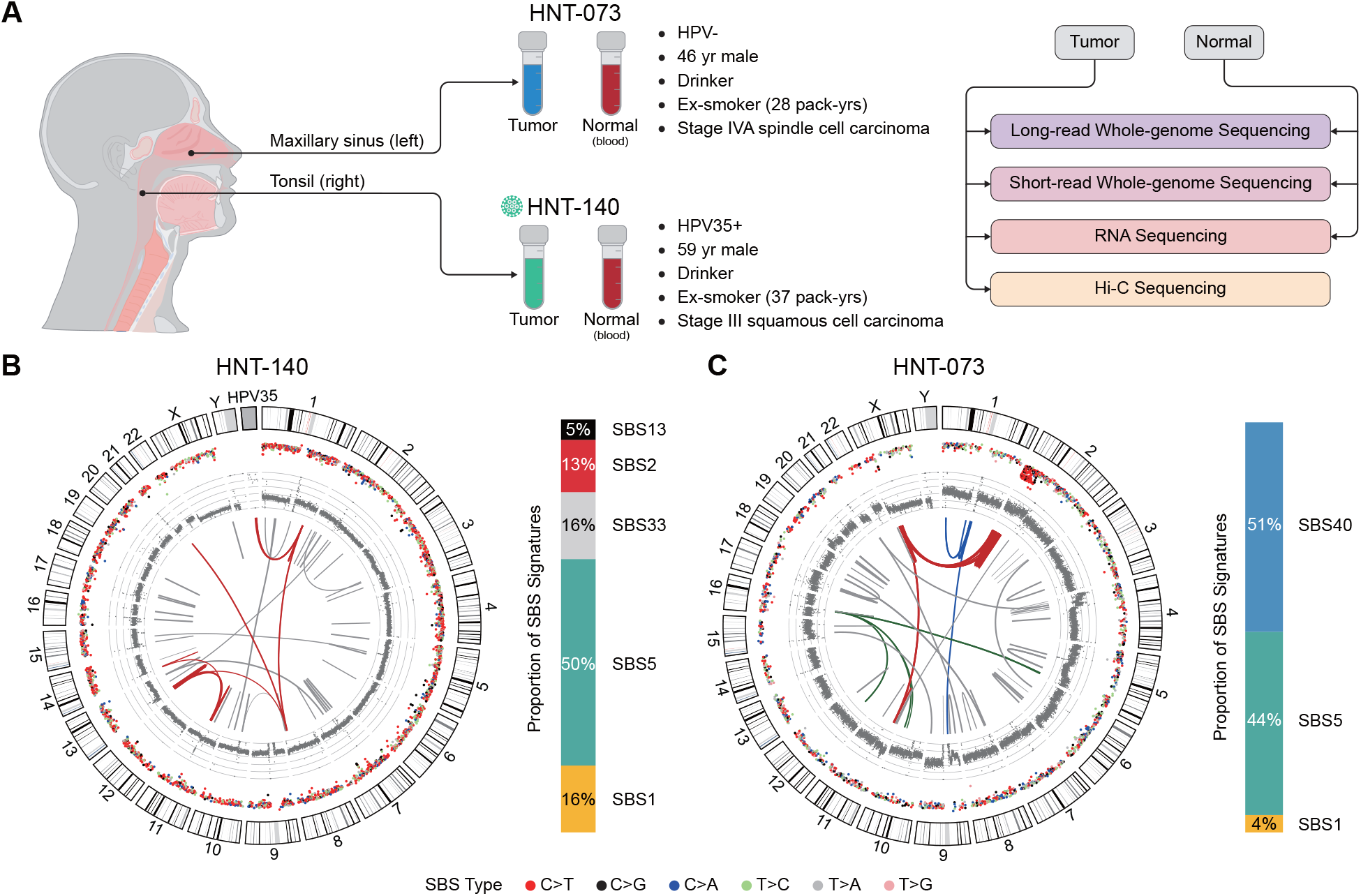
Overview of head and neck cancer patients, experimental design, genome-wide characteristics. (A) Patient characteristics and datasets generated in this study. (B-C) Genome-wide characteristics of HNT-140 and HNT-073. Starting from the outermost tracks in the circos plots: hg19 karyotype cytobands, intermutation distances (log_10_) of SBSs, sequencing depths (gray), and somatic structural variants (arcs). Clustered SVs are shown in different colors. The red arcs in HNT-140 represent amplifications that originate from the HPV integration site. The red arcs in HNT-073 represent the multichromosomal chromothripsis originating from chromosome 2. The blue and green arcs in HNT-073 represent SV clusters that did not result in any classification. The gray arcs in both samples represent SVs that were not clustered. The stacked bar plots to the right of each circus plot shows the SBS mutational signatures identified in each sample. (B) HNT-140 shows a multi-chromosomal chained SVs (red) that include HPV35 integration and APOBEC-associated signatures. (C) HNT-073 also shows a multi-chromosomal chromothripsis with *kataegis* and a homozygous deletion of *CDKN2A*.

HNT-140 is a case with HPV35 viral genome integration on chromosome 1q of the tumor genome (Fig 1B, Fig EV2). The integration breakpoints flanked translocations to 8q24, 13q13, and 11q13, suggesting that these events are chained. Although there were no driver sequence mutations, the APOBEC mutation signatures SBS2 and SBS13 were present in 18% of somatic mutations (Table EV7), indicating dysregulated activities of APOBEC cytidine deaminases found in HPV-positive HNC (Henderson *et al*, 2014). HNT-073 was a case with inter-chromosomal chromothripsis comprised of 180 SVs in 2p, 11p and Xp, with localized hypermutations, *kataegis* (Fig 1C).

First, by taking advantage of LR WGS, we developed a novel analytic method to capture simultaneous occurrence of two types of mutations in the same single molecule. We identified SBSs that only present in long reads supporting SV and referred to as ‘post-SV SBS’ (Fig 2A, Fig EV4 and Table EV8). Of the 570 SBSs identified as post-SV SBSs in 2p in HNT-073, 414 were in the chromothripsis cluster indicating *kataegis.* Approximately 40% and 9% of the SBSs co-occurring with the chromothripsis and the non-chromothripsis clusters were attributable to APOBEC activity, respectively (Fig 2B and Fig EV5). The evidence of APOBEC activity on generation of the post-chromothripsis *kataegis* is reinforced by the remarkably lower proportion of the aging signature SBS5 - 17% and 75% in the chromothripsis and non-chromothripsis SBSs, respectively. Double base substitutions of post-chromothripsis also indicated association with APOBEC activity (Fig EV6). This finding is consistent with the underlying mechanism of chromothripsis and *kataegis,* commonly associated with AID/APOBEC activity in multiple cancer types (Davis *et al*, 2014; Nik-Zainal *et al*, 2012).

**Figure 2.**
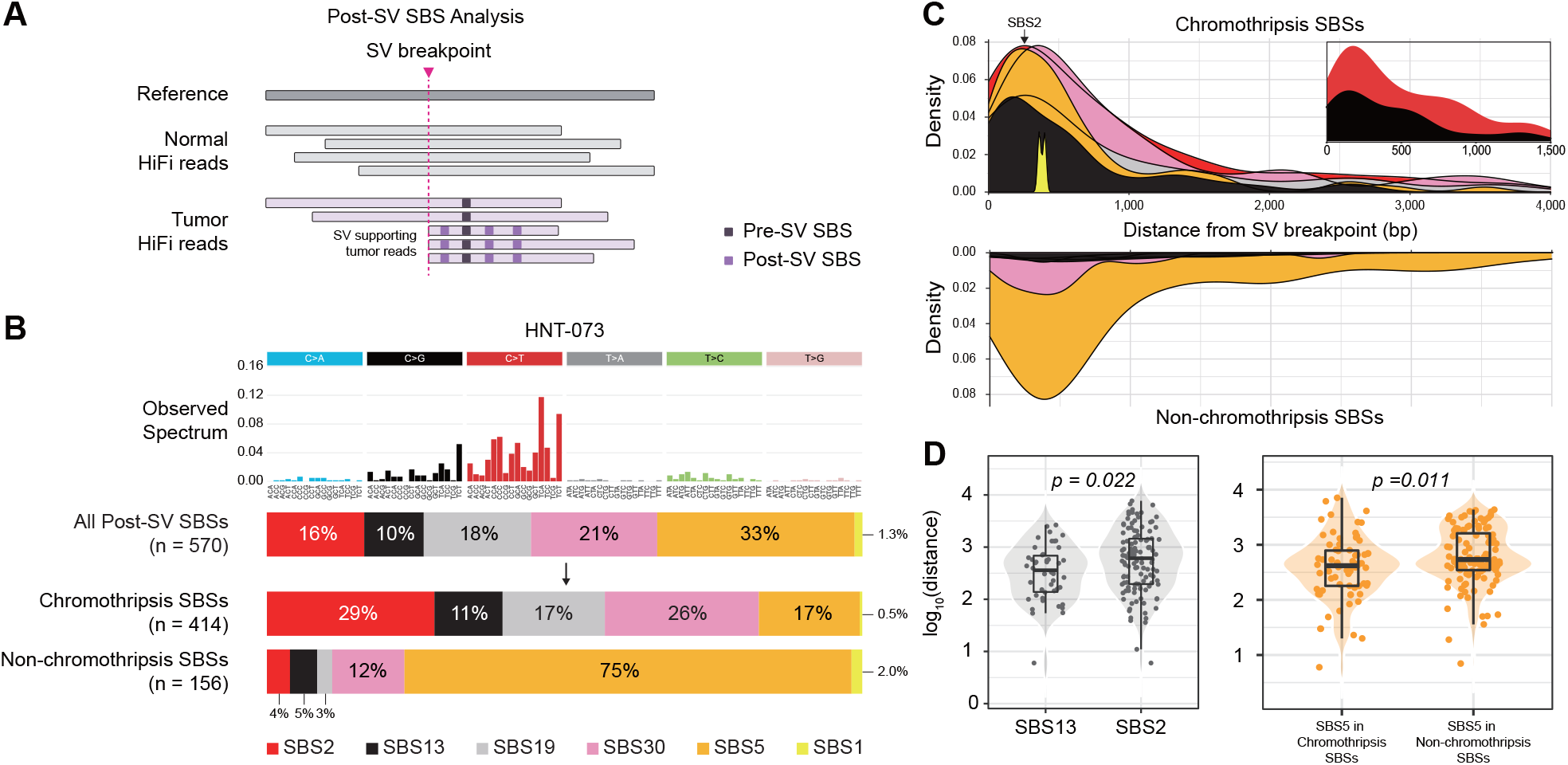
Post-SV mutational signature patterns identified using long-read sequencing. (A) A conceptual illustration of post-SV SBS analysis. (B) The observed spectrum of post-SV SBSs of HNT-073 and mutational signature proportions of the post-SV SBSs. Signature weights of all post-SV SBSs, chromothripsis-associated SBSs *(kataegis),* and non-chromothripsis-associated SBSs are shown in horizontal stacked bar plots. (C) The density of each signature between the two groups (chromothripsis-associated SBSs and non-chromothripsis SBSs) at varying genomic distances from the SV breakpoint. (D) The left boxplot shows genomic distances (log_10_) of SBS13 and SBS2 signatures from post-SV chromothripsis SBSs (Wilcox test *p* = 0.022). The right boxplot shows genomic distances (log_10_) of SBS5 signature variants in the chromothripsis cluster and the non-chromothripsis cluster from SV breakpoints. The chromothripsis cluster SBSs are closer to the SV breakpoints compared to those of the non-chromothripsis cluster SBSs (Wilcox test *p* = 0.011).

Using MutaliskR, we decomposed the variant-level mutational signatures of the post-SV SBSs in HNT-073. The density of SBS2 and SBS13 peaked at 297-bp and 176-bp, respectively, in the chromothripsis SBSs (Fig 2C). SBS13 mutations were found to be significantly closer to the SV breakpoints (Wilcox test *p* = 0.022; Fig 2D). This suggests potentially different APOBEC enzymatic activity roles at chromothripsis breakpoints. The genomic distances of SBS5-dominated post-SV SBSs from SV breakpoints were also significantly smaller (p = 0.011) for the SBSs of the chromothripsis cluster compared to those of the non-chromothripsis cluster (Fig 2D). Our analytic approach may reveal the unknown association of mutation patterns with mechanisms of SV generation in a large set of samples.

We next integrated the generated multi-modal sequencing data to infer the tumorigenesis history by considering the somatic SVs. First, in HNT-140, we reasoned that persistent viral infection preceded tumorigenesis as the HPV35 integration was clonal (SV allele frequency = 0.32) and the viral oncogenes E6 and E7 expression was intact. The host-viral genome interaction was also observed in the 3D genome conformation analysis (Fig 3A). HPV types 16 and 18 integrants flanking focal genomic amplifications are known to be evidence of looping of host-viral integrated genomic segments (Akagi *et al*, 2014). Based on *de novo* assembly of LR WGS, we found that type 35 also constructs an HPV integration mediated DNA looping and local amplification in primary HNC patient genome, leading to genomic instability (Table EV9).

**Figure 3.**
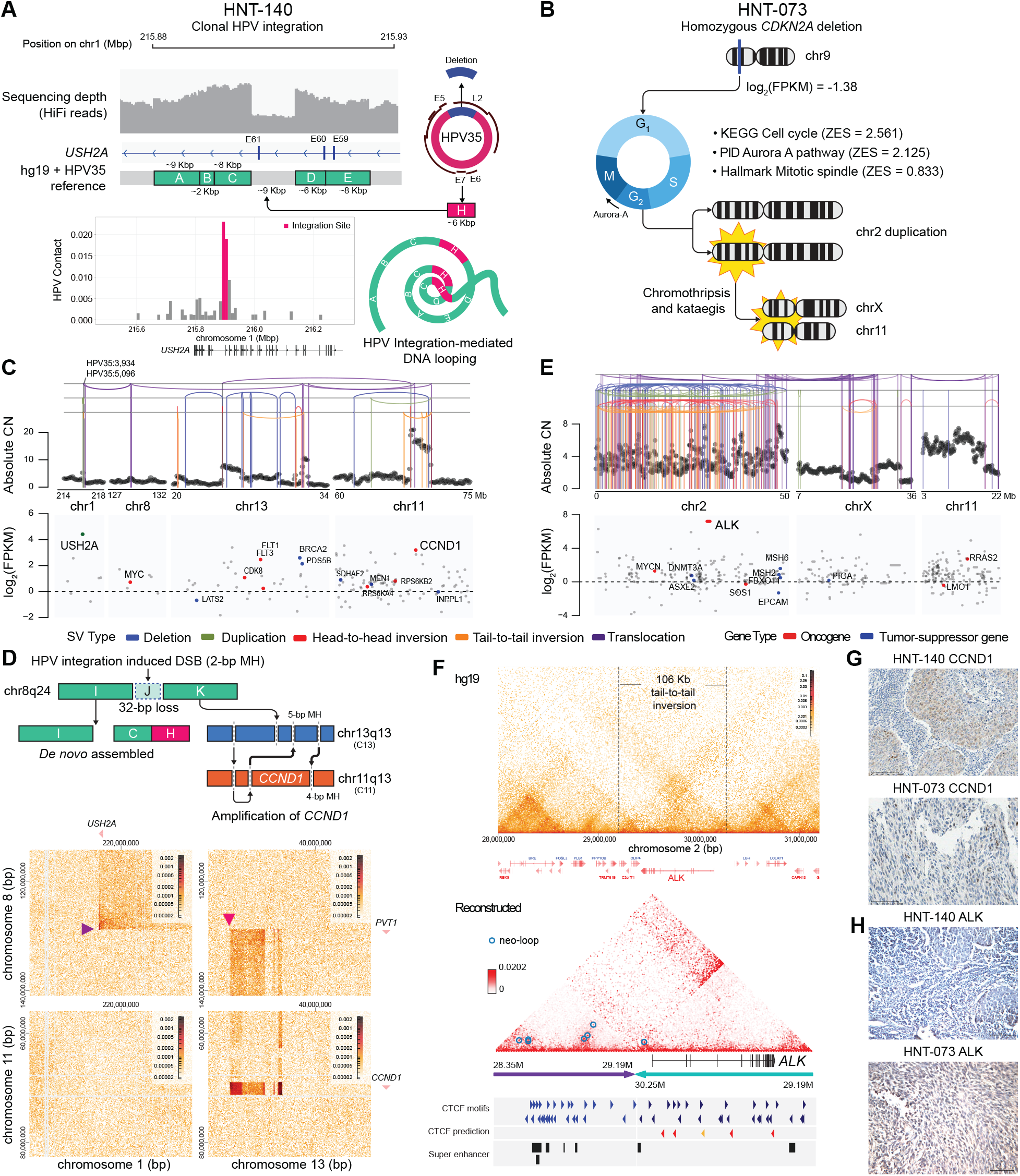
Inferring evolutionary history of head and neck carcinogenesis. (A-B) Early clonal oncogenic events of the HPV-positive HNT-140 and the HPV-negative HNT-073 cases are shown. (A) The genome structure and chromatin interactions around the HPV integration site and the integration-mediated DNA looping in HNT-140. (B) The cascade of tumor initiating events in HNT-073. (C) The top panels show SV breakpoints across HPV-integration-mediated complex rearrangements in HNT-140. The bottom panels show mRNA log_2_ fold changes in expression in the respective regions. (D) Schematic illustration of the HPV integration site flanking SV cluster resolved via *de novo* assembly. The chromatin interaction (bottom) plot shows evidence of concordant translocation supported by the two assembled contigs (I-B-C-H and K-C13-C11). (E) Top panels: SV breakpoints across chromothripsis in HNT-073. Bottom panels: mRNA log_2_ fold changes in expression in the respective regions. (F) The top panel shows the chromatin interaction plot showing *ALK* in the reference genome (hg19) coordinates. The bottom panel shows the augmented chromatin interaction data with the somatic inversion incorporated as well as the CTCF motifs, CTCF predictions, and presence of super enhancers (SEdb) in the augmented genomic segment. (G-H) Immunohistochemistry staining of HNT-140 and HNT-073 tumor samples for CCND1 and ALK.

HNT-073 harbored a 2.8 Mb homozygous deletion in 9p21 (SV allele frequency = 0.43 and log-2 copy-number ratio = −1.5) (Fig EV3). The deletion included the tumor suppressor *CDKN2A,* one of major driver gene in HNC (Fig 3B) (Cancer Genome Atlas, 2015). The single-sample gene-set enrichment analysis of HNT-073 showed overexpression of cell cycle pathways including Aurora-A and mitotic spindle pathways (Table EV10). Silencing of *CDKN2A* has been demonstrated to disrupt cell cycle by upregulating *AURKA* and *CCND* (Helias-Rodzewicz *et al*, 2018). Dysregulated *AURKA* is known to contribute to generation of mitotic DNA damage (Barr & Gergely, 2007) and severe genomic instability in cancer (Torchia *et al*, 2009). The chromothripsis on 2p, and amplifications of multiple chromosomes (Table EV11) in HNT-073 are consistent with the previously documented roles of *AURKA* in oncogenesis.

We next profiled differentially expressed genes near the breakpoints of the inter-chromosomal SV cluster associated with the HPV integration sites in HNT-140 and the chromothripsis cluster in HNT-073 (Table EV14). In HNT-140, the HPV integrant involving *USH2A* extended to an intronic region of the *PVT1* oncogene in 8q24, 13q13 and 11q13 (Fig 3C upper panel). Notably, the oncogene *CCND1* in 11q13 was amplified and overexpressed in HNT-140 with a log_2_ fold change (FC) of 3.20 (Fig 3C lower panel). Furthermore, *de novo* assembly and Hi-C interaction data further resolved this HPV-mediated chained SVs. We found that the somatic rearrangement is comprised of two clusters: a set of contigs with HPV fragments and the remaining contigs with a high level of microhomology at junctions near *CCND1* and *BRCA1* (Fig 3D). Our results suggest that HPV integration triggers a series of SV evolution that disrupt oncogenes or tumor suppressors to provide a selective advantage in HNC in line with previous hypothesis (Jeon *et al*, 1995; Moody & Laimins, 2010).

In HNT-073, in the regions of chromothripsis involving hundreds of genomic rearrangements, the oncogene *ALK* was significantly overexpressed (log_2_FC = 7.21, Fig 3E). We found a somatic inversion involving *ALK,* supported by a butterfly pattern of the 3D genome contacts (Fig 3F) (Tigano *et al*, 2021). Reconstruction of the 3D genome interaction of the *ALK* inversion region identified several neoloops, including one immediately upstream of the inverted *ALK* (Fig 3F, Table EV12-EV13). This region also included a super enhancer found in melanoma (COLO679) based on SEdb (Jiang *et al*, 2019), suggesting a synergistic effect among the oncogenic regulatory elements, the neo-loop, and the super-enhancer, altogether strongly activating the downstream *ALK* expression.

The protein expression levels of CCND1 and ALK in the two HNC patient samples were quantified using immunohistochemistry and concordant with the RNA-seq estimates (Fig 3G-H). In the TCGA-HNC study, 76 of 243 (31%) of HPV-negative tumors have *CCND1* amplification, but this is rare in patients with HPV positivity (1 of 36 cases) (Cancer Genome Atlas, 2015). Similarly, *ALK* overexpression was found in fewer than 5% of patients with HPV-negative TCGA-HNC (7 of 279 cases, 2.5%) (Cancer Genome Atlas, 2015). Although rare, CCND1 and ALK are known as predictive or prognostic factors in many cancer types (Feng *et al*, 2011; Kalish *et al*, 2004; Lee *et al*, 2014; Lin *et al*, 2017; Ouyang *et al*, 2018) and also have therapeutic potential (Aubry *et al*, 2019; Gougis *et al*, 2019; Holla *et al*, 2017; Michel *et al*, 2016). Given that cetuximab, which targets EGFR, a member of receptor tyrosine kinase family, is currently FDA-approved for HNC, ALK could be a candidate for future therapeutic choices.

Despite a limited number of cases, integrative profiling of complex genomic rearrangements using LR WGS, Hi-C sequencing, and RNA-seq results in deciphering rare driver SV events in HNC patients that lack driver SBS events. We expect such comprehensive analyses to aid discovery of rare and novel targetable driver mutations in other tumor types and inform personalized therapeutic strategies going forward.

## MATERIALS AND METHODS

### Sample Collection

Clinicopathological data, tumor and matched normal (blood) samples of head and neck cancer patients were collected with the Institutional Review Board (IRB) approval and informed consent at the National Cancer Center Korea.

### Ethics approval and consent to participate

Primary head and neck tumor tissues from the patients were previously banked at the National Cancer Center Korea with informed consent in accordance with the Declaration of Helsinki and IRB-approved research protocols (IRB No.NCCNCS08200).

### HPV Status

To determine the presence of HPV, we used a total of 10 mL of purified tumor DNA was used in each HPV PCR. Genotyping was performed using a PCR-based HPV DNA Chip (Green Cross, Gyeonggi, Korea) (Park *et al*, 2012).

### High-fidelity (HiFi) Circular Consensus Sequencing Library Preparation and Data Preprocessing

Fresh frozen head and neck tumor tissues were ground using the Qiagen TissueLyser II. The DNA from the tumor tissues and the matched normal (blood) samples were extracted using a DNeasy Blood & Tissue Kit (Catalog #69504, Qiagen). 8 μg of input genomic DNA was used for the HiFi library preparation. For genomic DNA where the size range was less than 17 kb, the Femto Pulse System (Agilent) was used to determine the actual size distribution. We sheared the genomic DNA with the Megaruptor^®^ 3 (Diagenode) and purified them using AMPure PB magnetic beads (Pacific Biosciences) if the genomic fragment size was greater than 40 kb. We prepared a total of 10 μL HiFi library using the PacBio SMRTbell Express ^®^ Template Prep Kit 2.0. Using Sequel II Bind Kit 2.2 and Int Ctrl 1.0, we annealed the prepared SMRTbell templates. We used Sequel II Sequencing Kit 2.0 along with SMRT cells 8M Tray to perform the sequencing. For each SMRT cell, 30-hr movies were captured on the Sequel sequencing platform by Macrogen Inc. (Seoul, Korea), generating subreads on the Pacific Biosciences Sequel II system. Using subreads as input, circular consensus sequences (CCS) with a minimum of 3 passes and minimum predicted accuracy of 0.99 were generated using CCS (v.4.2.0) tool, yielding high quality CCS results. The resulting raw CCS reads were 60.8 and 52.5 Gbp in blood and 65.8 and 61.4 Gbp in tumor samples in HNT-140 and HNT-073, respectively.

### Hi-C Sequencing Library Preparation

The patient tumor sample was removed from the liquid nitrogen storage and weighed while still frozen. Using a mortar and pestle on a bed of dry ice, the samples were pulverized while frozen until they formed a fine powder. The pulverized tissue was transferred to a 15mL tube containing 10mL of 1X DPBS and fixed using 2% formaldehyde for 10 minutes. The samples were quenched with 0.2M glycine for 5 minutes. The tissue was washed twice in 1X DPBS. After removing of the supernatant, the pellets were frozen at −80°C until further processing. Starting Hi-C experiments, the pellets were thawed and resuspended in 3mL of tissue lysis buffer (10mM Tris-HCl pH 8.0, 5mM CaCl2, 3mM MgAc, 2mM EDTA, 0.2mM EGTA, 1mM DTT, 943 0.1mM PMSF, 1X Complete Protease Inhibitors). We then transferred the samples to an M tube and subsequently ran them on the “Protein M-tube” program using a gentleMACs tissue dissociator (Miltenyi). After the dissociation, an additional 3mL of tissue lysis buffer with 0.4% Triton X 100 was added to the sample and the solution was passed through a 40μM strainer.

The tube and cell strainer were washed with an additional 2mL of 0.2% tissue lysis buffer with 0.2% Triton X-100. The sample was centrifuged and washed with an additional 1mL of tissue lysis buffer with 0.2% Triton X-100. For the tissue samples processed as described above, the digestion and Hi-C library preparation proceeded in a similar manner as based on the in situ Hi-C protocol (Rao *et al*, 2014). The pellet was suspended again in 50μL of 0.5% SDS, incubated for 10 minutes at 62°C. 145μL of water and 25μL of 10% Triton X-100 were added to quench the SDS for 15 minutes at 37°C. 25μL of 10x NEB Buffer 2 was added to the samples. Cells were digested with 500U of MboI restriction enzyme (NEB) in NEB Buffer 2. DNA ends were filled in with dNTPs, including biotin-14-dATP (Jena), using Klenow polymerase (NEB). Chromatin ends were ligated using T4 DNA ligase. DNA was then purified and sheared on a Covaris M-series ultrasonicator. Biotinylated fragments were purified using the My T1 Streptavidin coated beads (Life Technologies) and subject to on-bead library preparation as described previously (Rao *et al.,* 2014).

### Illumina Short-read Sequencing Library Preparation

The input DNA materials were extracted and processed in the same way as described above (PacBio HiFi CCS Library Preparation and Data Preprocessing) for the Illumina short-read whole-genome sequencing. The samples were prepared according to the Illumina TruSeq Nano DNA library preparation guide. 100 ng of DNA was sonicated with a Covaris S220 Focused-ultrasonicator, and 151-bp paired-end libraries were constructed.

RNA extraction of patient’s tumor and matched adjacent normal tissue samples was performed using RNeasy Mini Kit (Catalog #74104, Qiagen). We calculated the total RNA concentration using Quant-IT RiboGreen (Invitrogen) and determined the DV200 (% of RNA fragments >200 bp) value using TapeStation RNA screentape (Agilent). A total of 100 ng of RNA was used as input to the mRNA sequencing library prep kit (TruSeq RNA Access Library Prep Kit by Illumina, San Diego, CA, USA) according to the protocols of the manufacturer. Finally, the 101-bp paired-end libraries were prepared and used for sequencing.

### Sequencing Statistics

Short-read WGS was performed on Illumina HiSeq X Ten instruments with read lengths of 2 × 151 bp, yielding approximately 1.4 billion mapped reads from the tumor samples (mean coverage of 68X) and 770 million mapped reads from the normal samples (mean coverage of 37X). mRNA sequencing (RNA-seq) was performed on Illumina HiSeq 2500 instruments (Illumina, Inc., San Diego, CA, USA) with read lengths of 2 × 101 bp, generating approximately 51 million aligned reads from the tumor samples and 24 million aligned reads from the normal samples (Supplementary Table 2). Hi-C libraries were sequenced using the Illumina Novaseq S4 as paired-end 150-bp reads and approximately 1.2 billion reads were aligned to the reference genome. Diagram of following analysis workflow is visualized in Fig EV7.

### Long-read Whole-genome Sequencing Analysis Workflow

The long-read (LR) genome sequencing reads were aligned to hg19 using minimap2 (Li, 2018) (v2.20). The reference-aligned long-reads were used to call single-nucleotide variants (SNVs) and small insertions and deletions (INDELs) using GATK4 Mutect2 (McKenna *et al*, 2010) and DeepVariant (Poplin *et al*, 2018). Structural variants were detected using Sniffles (Sedlazeck *et al*, 2018) and pbsv (https://github.com/PacificBiosciences/pbsv). Taking advantage of the long-reads, we also performed *de novo* assembly using hifiasm (Cheng *et al*, 2021). The assembled contigs underwent variant calling using *paftools* (distributed with Minimap2). The raw variant callsets were refined based on the number of supporting reads, software-specific filter parameters and quality scores wherever possible. Germline calls were screened and filtered out based on calls identified in the HG002 normal sample (Nurk *et al*, 2022). The refined sets of SNV, INDEL and SVs were subsequently annotated using ANNOVAR (Wang *et al*, 2010).

### Short-read Whole-genome Sequencing Analysis Workflow

The short-read (SR) genome sequencing reads were aligned to hg19 using bwa-mem (Li, 2013). Local realignment was performed using GATK3 (McKenna *et al.,* 2010). We used Picard (http://broadinstitute.github.io/picard) to mark PCR duplicate reads, after which the base scores were recalibrated using GATK3. The postprocessed reference-aligned reads were then used for SNV and INDEL calling with GATK4 Mutect2. The somatic copy number alterations were identified using Sequenza (Favero *et al*, 2015) with the matched normal as the control sample. Structural variants were detected using Delly2 (Rausch *et al*, 2012). SV calls were processed and classified according to the approaches described by Lee et al. (Lee *et al.,* 2019).

### Comparison and Integration of LR and SR WGS Variant Callsets

Given the limitations of the low coverage and low tumor purity of the LR WGS dataset, we sought to generate a harmonized callset by comparing and integrating the somatic callsets (SNVs, INDELS, and SVs) obtained from the SR and LR datasets. We note that SVs and somatic mutations exclusively found in the high-coverage SR WGS dataset had significantly lower cancer cell fractions (CCF) compared to the LR WGS dataset (Fig EV9).

### Short-read RNA Sequencing Analysis Workflow

The abundance of hg19 transcripts was quantified using RSEM (Li & Dewey, 2011) for both the tumor and the normal samples. Significantly enriched gene sets were identified by ssGSEA (Yi *et al*, 2020) based on Hallmark, PID, and KEGG databases through GenePattern cloud (Reich *et al*, 2006).

### Hi-C Sequencing Analysis Workflow

We followed the HOMER pipeline (Heinz *et al*, 2010) for the general processing of the Hi-C sequencing reads. The Hi-C sequencing reads of the tumor samples were initially trimmed using HOMER and then aligned to hg19 using bowtie2 (Langmead & Salzberg, 2012). A/B compartments were called using HOMER. The HOMER generated HIC files were converted to COOL files and MCOOL files using HiCExplorer (Ramirez *et al*, 2018) and Juicer (Durand *et al*, 2016), respectively. Putative somatic translocation calls from the SR and LR WGS datasets were verified using HiGlass (Kerpedjiev *et al*, 2018) by visual inspection.

### Identification of Neoloops and Expressed Structural Variants

High-confidence SV callsets along with the Hi-C sequencing data were used to identify neoloops using NeoLoopFinder (Wang *et al*, 2021). Predicted neoloops were visually inspected using the visualization module in NeoLoopFinder. Super enhancers were referenced from SEdb (Jiang *et al.,* 2019) for the regions flanking the predicted neoloops.

### Post-SV SBS Variant-level Signature Analysis

We developed MutaliskR (https://github.com/Honglab-Research/MutaliskR), which extends our previously published maximum likelihood estimation method of mutational signature analysis (Lee *et al*, 2018) to double-base substitutions (DBS) and small insertions and deletions (INDELs). It assigns variant-level signatures as previously described by Letouze et al. (Letouze *et al*, 2017). By taking advantage of long reads, we also developed a novel analytical approach that investigates variant-level mutational signatures of SBSs flanking SV breakpoints. We extracted reads that support each SV and performed mutation calling using Mutect2 (Cibulskis *et al*, 2013) using the reads. With manual IGV inspection, mutations only shown in SV-supporting reads and supported by two or more reads were further investigated. SVs with more than two mutations co-occurred were used in the post-SV SBS analysis. Variant-level signature analysis was performed on the post-SV SBSs using MutaliskR.

### Immunohistochemistry

We evaluated CCND1 and ALK protein expression using immunohistochemistry assays. The paraffin blocks of the tumor samples were sectioned and marked on standard hematoxylin/eosin. Slides were stained with the rabbit monoclonal antibody for ALK (clone D5F3, the Cell Signaling Technology, USA) and Cyclin D1 (P2D11F11, Novocastra Laboratories Ltd., Newcastle, UK) per the manufacturer’s instructions and visualized with DAB chromogen (Dako, USA).

## Data Availability

HiFi long read whole genome data: Sequence Read Archive (SRA) SUB11971936 (https://www.ncbi.nlm.nih.gov/sra)

## ACKNOWLEDGMENTS

This work was supported by the National Research Foundation of Korea (NRF) grant funded by the Korean government (Ministry of Science and ICT) (No. 2020R1C1C1011379 for J.C., 2021M3H9A2097227 and 2022R1A2C3008162 for D.H., and 2020R1A2C2005091 for Y.S.J.), the National Cancer Center (1810862-4 and 2031130-3 for Y.S.J) and the Catholic Medical Center Research Foundation made in the program year of 2020 (for D.H.)

